# Expansion of chromosome F heterochromatin in parthenogenetic *Drosophila mercatorum*

**DOI:** 10.1101/2025.06.22.660940

**Authors:** A. L. Sperling, D. K. Fabian, D. M. Glover

**Affiliations:** University of Cambridge; Department of Genetics, University of Cambridge, Cambridge CB23EH, UK; Division of Biology and Biological Engineering, California Institute of Technology; Pasadena, California 91125, USA

**Author notes:** Corresponding authors. A. L. Sperling D. M. Glover.

**Keywords:** Parthenogenesis, Chromosome Inversions, *Drosophila*, Heterochromatin, Repetitive DNA

## Abstract

The dramatic transition to parthenogenetic reproduction is often accompanied by similarly dramatic changes in genome organization. Genomic changes may precede or succeed the onset of parthenogenesis and may directly or indirectly contribute to the success of this reproductive transition and fitness of parthenogenetic animals. To gain greater understanding of genomic changes accompanying the transition to parthenogenetic reproduction, we have characterized genomic differences between a sexually reproducing and parthenogenetic strain of *Drosophila mercatorum*. This revealed three large (2.6-9 Mbp) inversions present on Muller element B and a 30-fold length expansion of heterochromatin on Muller element F. Comparisons of these genomic changes and parthenogenetic ability in 18 *D. mercatorum* strains collected in South America, North America, Hawaii, and Africa indicates no clear correlation of these genomic changes with geographical origin. However, these changes are minimal in strains collected in Brazil suggesting this is the origin for the ancestral strain and that both chromosome inversions and increased heterochromatin have likely preceded the transition to parthenogenetic reproduction. Parthenogenetic ability correlates most strongly with expansion of chromosomal element F. We speculate that this increased heterochromatic environment of genes on element F influences gene expression to either enhance parthenogenesis directly or redirect the activity of another factor contributing to it, in a manner analogous to the known influence of heterochromatin on gene activity in position effect variegation.

## Introduction

Parthenogenesis has been proposed to be an adaptive trait that would be beneficial when colonizing new habitats or when at low population density (Kearney 2003; Chang, et al. 2014; Esposito, et al. 2024). This may be particularly true for the diverse *Drosophila* genus, of which an estimated 76% of species are capable of some, usually low, degree of parthenogenesis (Sperling and Glover 2023a). Parthenogenetic capability also appears linked to the geographical collection points for many different species pointing towards an association with events aiding geographical isolation (Stalker 1954, 1964; Carson 1967; Templeton, et al. 1976; Matsuda, et al. 2009; Chang, et al. 2014). Differences in parthenogenetic ability between strains of *Drosophila* species have been most extensively studied in the repleta group, in particular in *Drosophila mercatorum* (Carson 1967; Carson, et al. 1969; Carson 1973; Templeton, et al. 1976; Templeton 1979; Sperling, et al. 2023). Indeed, parthenogenesis was first observed inadvertently in strains of *D. mercatorum* described as having only sterile males lacking a Y chromosome (Wharton 1942; reviewed in Sperling and Glover 2023a), a consequence of X chromosome non-disjunction. *D. mercatorum* strains exist that are capable of sexual reproduction, facultative parthenogenesis, or complete parthenogenesis making the species an excellent model in which to examine genome organisation associated with parthenogenesis and/or geographical isolation (Sperling, et al. 2023).

A molecular cause of parthenogenesis in *D. mercatorum* was recently uncovered by identifying genes that are differentially expressed in the germlines of and parthenogenetically reproducing strains and testing the consequences of changing their patterns of gene expression in *D. melanogaster* (Sperling, et al. 2023). This study identified three genes that could induce parthenogenetic reproduction in the non-parthenogenetic *D. melanogaster*: *Myc* on Muller element A (the X chromosome); *Desat2* on Muller element E (the 3R chromosome); and *polo* on Muller element D (the X chromosome 3L). *Myc* and *polo* can act as proto-oncogenes and have roles in regulating cell cycle progression and cell proliferation (Sunkel and Glover 1988; Grifoni and Bellosta 2015). The Muller elements A-F are equivalent to the six chromosome arms of the *Drosophila* karyotype (Muller 1940); the individual Muller elements from different *Drosophila* species carry the same complement of genes but have undergone a high degree of rearrangement as a result of multiple sequence inversions. *Desat2* is predicted to enable the synthesis of unsaturated fatty acids and is known to impart cold tolerance in *D. melanogaster* (Dallerac, et al. 2000; Takahashi, et al. 2001; Greenberg, et al. 2003). A decrease in *Desat2* expression has been credited with causing the global expansion of *D. melanogaster* (Greenberg, et al. 2003; Sinclair, et al. 2007). These genes work in concert to cause parthenogenesis yet are located on completely different chromosome arms. How then might parthenogenetic reproduction be selected, bearing in mind that many changes are likely to be required for the survival and adaptation of parthenogenetic animals?

At a gross level, substantial differences have been documented in the genomes of parthenogenetic and sexually reproducing animals, including polyploidy, aneuploidy, inversions and translocations, in addition to smaller changes to specific genes (Jaron, et al. 2020; Sperling and Glover 2023a; Sperling and Glover 2024). The selection pressures underlying the acquisition of this genome variability remain unknown in most animals. Like many other parthenogenetic animals, naturally parthenogenetic *D. mercatorum* also demonstrate a high degree of genome variability and display aneuploidy in their tissues (Sperling and Glover 2024). In addition to aneuploidy, naturally parthenogenetic *D. mercatorum* were also found to differ from a sexually reproducing strain by the presence of three large inversions on Muller element B and an expansion of heterochromatin on Muller element F (Sperling, et al. 2023). The cause or benefit of the additional inversions on element B and the expansion of heterochromatin on element F within the *D. mercatorum* species is unknown. Many chromosome inversions on different chromosome arms and a polymorphic element F were thought to have become fixed in populations that crossed the geographical barrier of the Andes in South America (Wharton 1942; Wasserman 1992). However, the relationship between geographical isolation and the incidence of parthenogenesis in unknown.

Naturally occurring heterochromatin expansion has not previously been linked to parthenogenesis, but it has been studied with respect to position-effect variegation (PEV) (reviewed in Liu, et al. 2020), transposable element (TE) regulation (Stamidis and Zylicz 2023), and increased lifespan (Larson, et al. 2012). However, there are very few examples of animals that have strains of the same species with dramatic or visible differences in the composition of their heterochromatin and so the consequences of such variation have received little attention. However, it is conceivable that the observed expansion of heterochromatin could influence the expression of genes on element F through position effect variegation and elsewhere in the genome through sequestration of heterochromatin protein 1 (HP1) (reviewed in Liu, et al. 2020). Since parthenogenesis can be induced in *D. melanogaster* by the change in expression of three genes in the absence of obvious genomic changes (Sperling, et al. 2023), it is likely that some distant regulator drives changes in gene expression in parthenogenetic eggs and changes to heterochromatin potentially offer such a route.

In contrast to dramatic increases in heterochromatin, chromosome inversions have long been documented to suppress recombination (Sturtevant 1917) and have been proposed to contribute to adaptation, speciation, and the divergence between sex chromosomes (Berdan, et al. 2023). Taxonomically widespread links have been established between inversions and the evolution of different mating systems, changes in social organization, environmental adaptation, and reproductive isolation (reviewed in Wellenreuther and Bernatchez 2018). Inversions have been most studied with respect to environmental adaptation (Kapun and Flatt 2019; Westram, et al. 2022; Berdan, et al. 2023). By preventing recombination between different alleles or discrete blocks of genes, inversions have been proposed to offer the maintenance of advantageous genetic changes or gene regulation (reviewed in Wellenreuther and Bernatchez 2018). Conversely, the prevention of recombination by inversions may prevent the removal of deleterious mutations or the recombination of different alleles to generate variations (discussed in Roesti, et al. 2022). Therefore, the selective value of inversions should be addressed in the context of physiological adaptation to different environments.

Elevated genetic divergence has been identified around the inversion breakpoints in a wide range of animal and plant species (*Drosophila,* Noor, et al. 2007; Kulathinal, et al. 2009; Lohse, et al. 2015; Poikela, et al. 2023; mosquitoes, Besansky, et al. 2003; Michel, et al. 2006; Cheng, et al. 2018; a butterfly, Joron, et al. 2011; shrews, Basset, et al. 2006; Yannic, et al. 2009; and sunflowers Rieseberg, et al. 1999; Barb, et al. 2014). Multiple adaptive consequences have been linked to inversions in *D. melanogaster* and *Drosophila pseudoobscura* (Dobzhansky 1947; Knibb, et al. 1981; Prevosti, et al. 1984; Prevosti, et al. 1985; Prevosti, et al. 1988; Kapun, et al. 2016; Nunez, et al. 2023) and inversions in flies adapted to high latitude increase stress resistance and longevity (Fabian, et al. 2012; Durmaz, et al. 2018) but effects of inversions on the specific expression of individual genes has yet to be shown. Moreover, the underlying mechanisms for the formation of genome inversions are also not fully understood. In the repleta group species, *Drosophila buzzatii*, inversions result from ectopic recombination at regions that are dense with transposable elements (Cáceres, et al. 1999). However, in most cases, the cause of inversions is not always evident from the genome sequence and the advantage they may impart is not clear.

Here, we wished to understand whether the genesis of specific chromosome inversions and expansion of heterochromatin precedes or is a consequence of parthenogenesis or selective pressures reflecting geographical isolation. To this end, we examined the karyotypes of 18 *D. mercatorum* strains collected in South America, North America, Hawaii, Africa, and from undocumented locations. We found that those strains capable of parthenogenesis showed an expansion of element F to a length similar to the other chromosome arms. Additionally, we identified inversions that were present in all strains capable of reproducing by parthenogenesis. The genomic changes that we describe are found in non-parthenogenetic strains, indicating that they alone do not cause parthenogenesis. However, the expansion of element F correlates strongly with parthenogenesis suggesting that this might well be a contributing factor.

## Results

*D. mercatorum* has the typical *Drosophila* karyotype consisting of 5 pairs of chromosome arms, known as Muller elements A-E, and a pair of small ‘dot’ chromosomes, Muller element F (Wharton 1942; DeSalle, et al. 1986). In the ancestral karyotype of the repleta species group the chromosome arms are all telocentric (Wasserman 1960). Whereas in *D. mercatorum*, there is a distinctive fusion between element B and element E (Wasserman 1963). Like other repleta group species, *D. mercatorum* likely originates from South America (Acurio 2024). It is thought to have migrated over the Andes, with karyotype polymorphisms, chromosome inversions and variation in the size of the dot chromosome becoming fixed in the population as a consequence of a founder effect (Wharton 1942; Wasserman 1960; Wasserman 1992). Our laboratory recently compared the genomes of a sexually reproducing and a parthenogenetic strain of *D. mercatorum* and identified three large inversions (9 Mbp for *In(EB)A*, 2.6 Mbp for *In(EB)B*, and 7.1 Mbp for *In(EB)C*) on element B in the DNA sequence and banding patterns of the giant salivary gland chromosomes of the parthenogenetic strain (Fig. 1A, B) (Sperling, et al. 2023). These inversions are not present on element B in the closely related *Drosophila repleta* (Wharton 1942), and thus the ancestral *D. mercatorum* element B is likely that of the sexually reproducing strain from Brazil. In addition to the inversions, element F of the sexually reproducing strain was significantly smaller in larval neuroblasts (on average 0.57 μm; n = 50, σ = 0.14, s.e. = 0.02) than in the parthenogenetic strain (15.67 μm; n = 52, σ = 3.35, s.e. = 0.46) (Fig. 1C), the latter being of similar size to the other telocentric chromosome arms. When the length was normalized over the total length of all chromosome arms there was a significant (*p*-value = 2.22 x 10^-16^) increase in element F chromosome length in the parthenogenetic strain (Fig. 1D). Since the sexually reproducing strain was South American in origin and the parthenogenetically reproducing strain was Hawaiian (Sperling, et al. 2023), this raises the question of whether these genomic differences were related to geographical separation and/or parthenogenetic ability.

**Figure 1:**
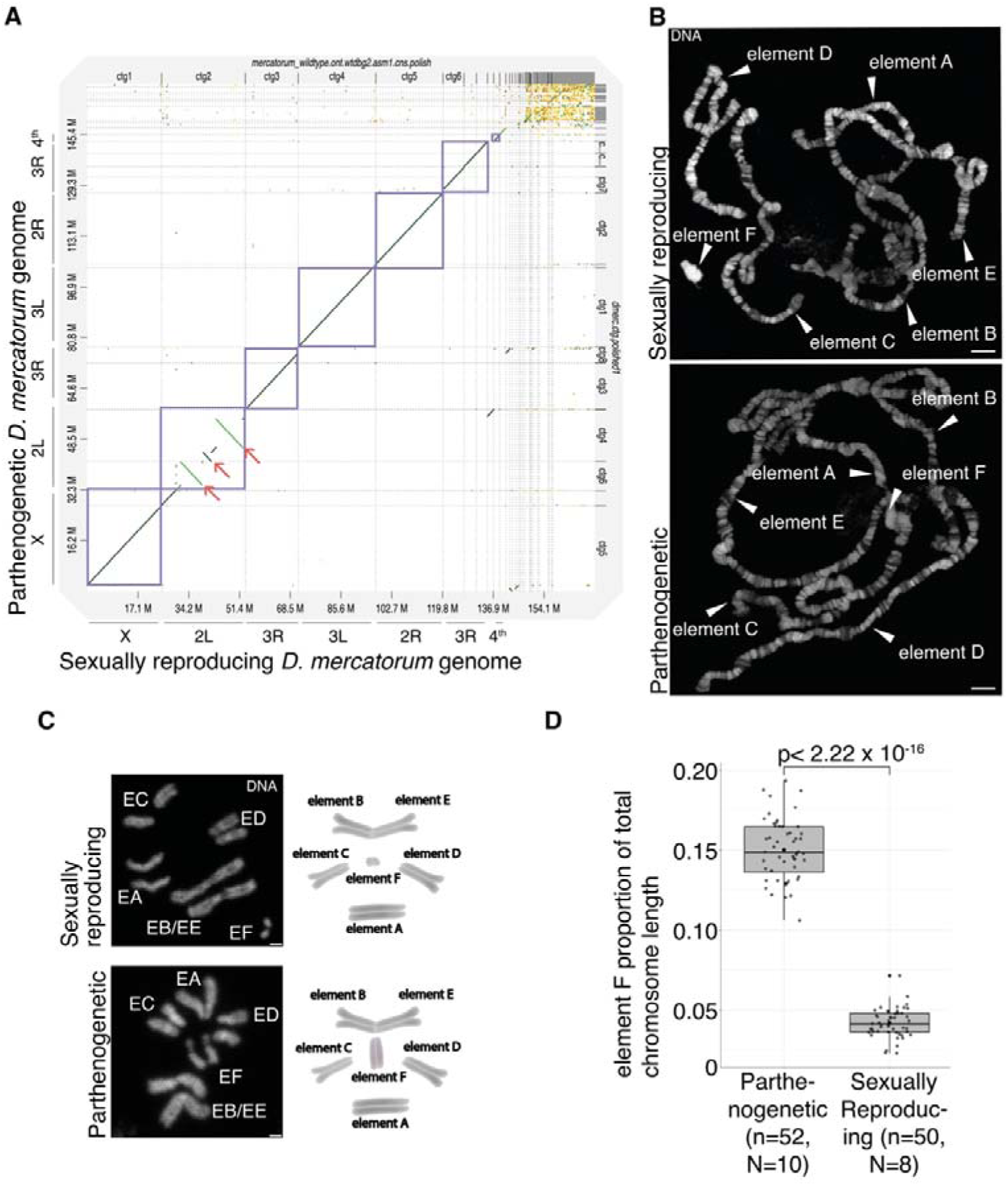
Genome differences between sexually reproducing and parthenogenetic *D. mercatorum*. **A**) Alignment of the genome assembly of a fully parthenogenetic *D. mercatorum* strain against the genome assembly of a sexually reproducing *D. mercatorum* strain. **B**) Polytene chromosome spreads from the salivary glands of parthenogenetic and sexually reproducing *D. mercatorum* 3^rd^ instar larvae in which the different chromosome arms are indicated. Scale bar, 10 μm. **C**) Mitotic chromosome spread from the brains of parthenogenetic and sexually reproducing *D. mercatorum* 3^rd^ instar larvae in which the chromosome arms are indicated. Preparations are stained with DAPI to reveal DNA. Scale bar, 1 μm. **D**) Box plot of element F chromosome length measured in confocal images using FIJI, normalized by the proportion of total chromosome length. *p*-value calculated with a t-test.

To gain more insight into the potential significance of these genomic differences, we first carefully examined the inversions on element B (Fig. 2A,B). Using Hybridization Chain Reaction (HCR) *in situ* with probes against genes on the main contigs that make up element B (Fig. 2C-D), we were able to relate the banding pattern of polytene chromosomes to the inversions. This enabled us to relate the three large-scale inversions predicted by the genome assembly to the physical organization of the polytene chromosomes in the parthenogenetic strain of *D. mercatorum* (Fig. 2E). We then manually annotated genes present in an arbitrary region of 100 kb around each of the breakpoints (Fig. S1A,B, Data Tables S1A-J) and determined if they were differentially expressed in the mature eggs of sexually reproducing and parthenogenetic strains to identify genes with potential to promote parthenogenesis (Data Table S1K). We identified four differentially expressed genes: Chronologically inappropriate morphogenesis (*chinmo*), encoding a transcription factor; *crossover suppressor on 2 of Manheim* (*c(2)M*), encoding a cohesin involved in synaptonemal complex assembly during meiosis; *squash* (*squ*), which encodes a piRNA pathway component involved in transposable element (TE) repression; and *CG17646* which encodes a transmembrane transporter. As our previous functional testing of the parthenogenetic capability of differentially expressed genes focussed primarily upon those regulating cell cycle or chromosome behaviour, we had previously only functionally tested *c(2)M* and found it not to be associated with parthenogenesis (Sperling, et al. 2023). Thus, it remains possible that *chinmo* (log_2_ fold change 3.1, padj = 1.1 x 10^-15^) could facilitate the development of parthenogenetic animals. Moreover, *squ* (log_2_ fold change 0.6, padj = 1.6 x 10^-2^) could be a candidate gene for stabilising transposable element (TE) activity in parthenogens.

**Figure 2:**
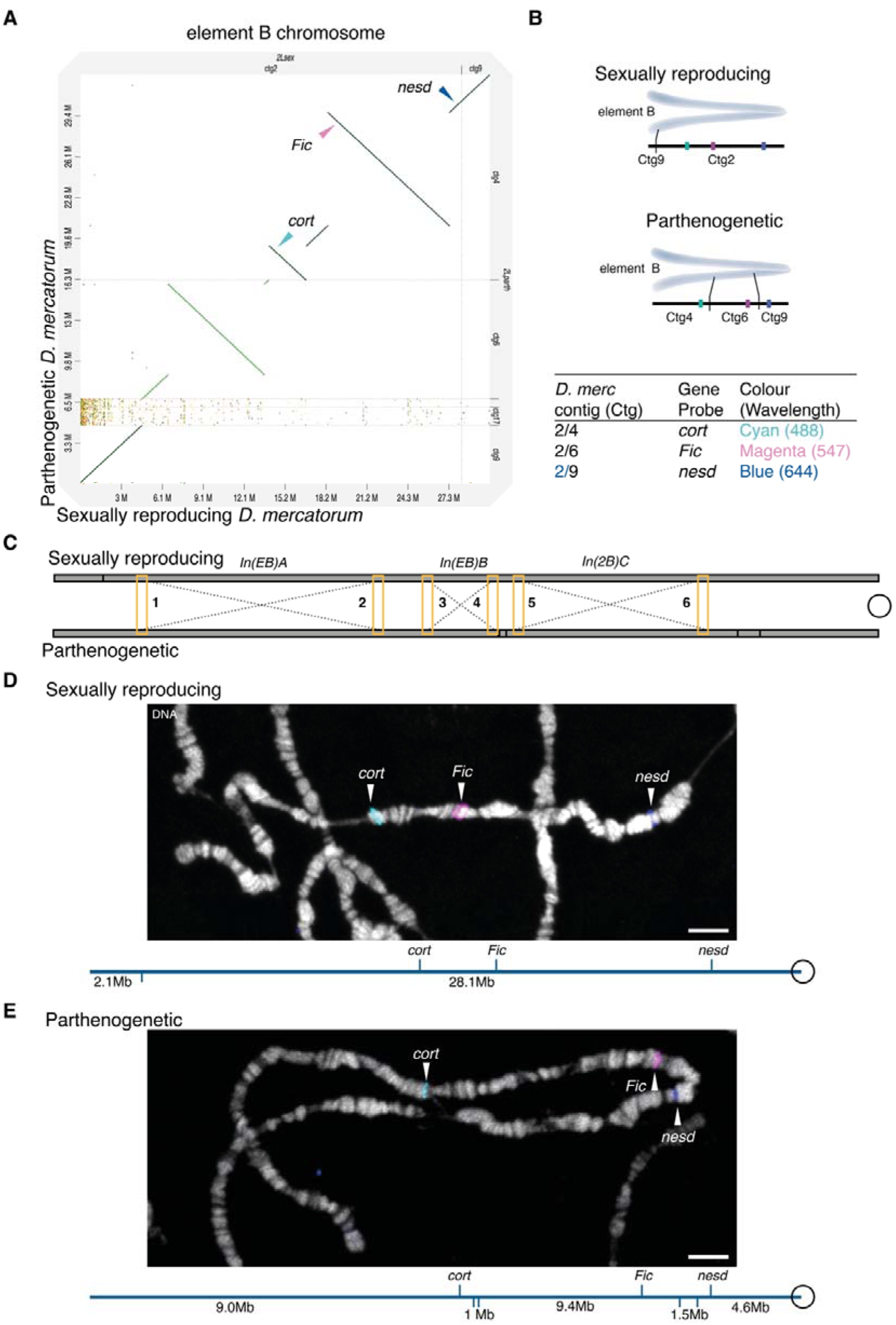
Differences in element B between sexually reproducing and parthenogenetic *D. mercatorum*. **A**) Alignment of the parthenogenetic against the sexually reproducing *D. mercatorum* element B with the approximate location of the three markers, *cort* (cyan), *Fic* (magenta), and *nesd* (blue), indicated. **B**) Predicted localization of select genes present on the contigs of element B for both sexually reproducing and parthenogenetic strains. **C**) Schematic of the inversions between the sexually reproducing and parthenogenetic *D. mercatorum* element B polytene chromosome arms. **D-E**) Polytene chromosome spread from the salivary glands of parthenogenetic and sexually reproducing *D. mercatorum* 3^rd^ instar larvae. *In situ* hybridization to select genes on the largest contigs to *cort* (cyan), *Fic* (magenta), and *nesd* (blue). The DNA (DAPI) is in monochrome. Scale bar, 10 μm. A schematic of the element B chromosome arm is given below the images.

We next searched for and found 5 differentially expressed genes within the entire 2.6 Mbp *In(EB)B* inversion (Fig. S1C): *CG31612* and CG4496, predicted to be involved in transcription; *Nha1,* a membrane transporter; *Nup107,* required in nuclear pore complex formation; and *Itgbn*, the Integrin betanu subunit. None of these were previously subjected to tests for their functional involvement in parthenogenesis (Sperling, et al. 2023). These newly identified genes are therefore candidate enhancers of the efficiency of parthenogenesis, survival, or adaptation. Our current method for testing the role in parthenogenesis in *D. melanogaster*, which recapitulates changes observed in *D. mercatorum* (Sperling, et al. 2023), does not allow us to investigate the role of additional genes. This is because we have been unable to test the combined effect of changing the expression of more than three genes in generating parthenogenesis in *D. melanogaster* due to decreases in viability (Data Table S1L). Therefore, an alternative approach will be needed to test the roles of new differentially expressed genes (see Discussion).

The other major change in the genome of the parthenogenetic *D. mercatorum* strain compared to its sexually reproducing counterpart is the size of element F (Fig. 3A,B). As element F of the parthenogenetic strain is 30-fold longer than that of the sexually reproducing strain, whereas the coding regions match along their length (Fig. 3C,D), the size expansion is likely due to non-coding repetitive DNA sequences that make up the non-assembled small contigs in the genomics data (Fig. S2). Simple repetitive DNA is the most likely candidate, since it forms the basis for establishing heterochromatin. Accordingly, the F elements of sexually reproducing and parthenogenetic strains are of similar size in polytene salivary gland chromosomes, which lack heterochromatin or repetitive DNA due to the absence of endoreduplication in these regions (Gall et al. 1971). Thus, the size difference in element F between the strains is likely due to the addition of satellite DNA or heterochromatin, as has been documented in other Hawaiian species of *Drosophila* (Craddock, et al. 2016) and accounting for the difficulty in assembling sequence contigs in this region. Moreover, genes present on the manually annotated element F corresponded to those on element F of the *D. melanogaster* reference genome (release 6) (Data Table S1M). We found five of these genes to be differentially expressed between the parthenogenetic and sexually reproducing transcriptome datasets (Sperling, et al. 2023): *apolpp*, *CG33978*, *Thd1*, *NfI*, and *mGluR*, possibly as a result of the changed organization of heterochromatin (see Discussion). None of these genes has been previously implicated in parthenogenesis (Data Table S1N).

**Figure 3:**
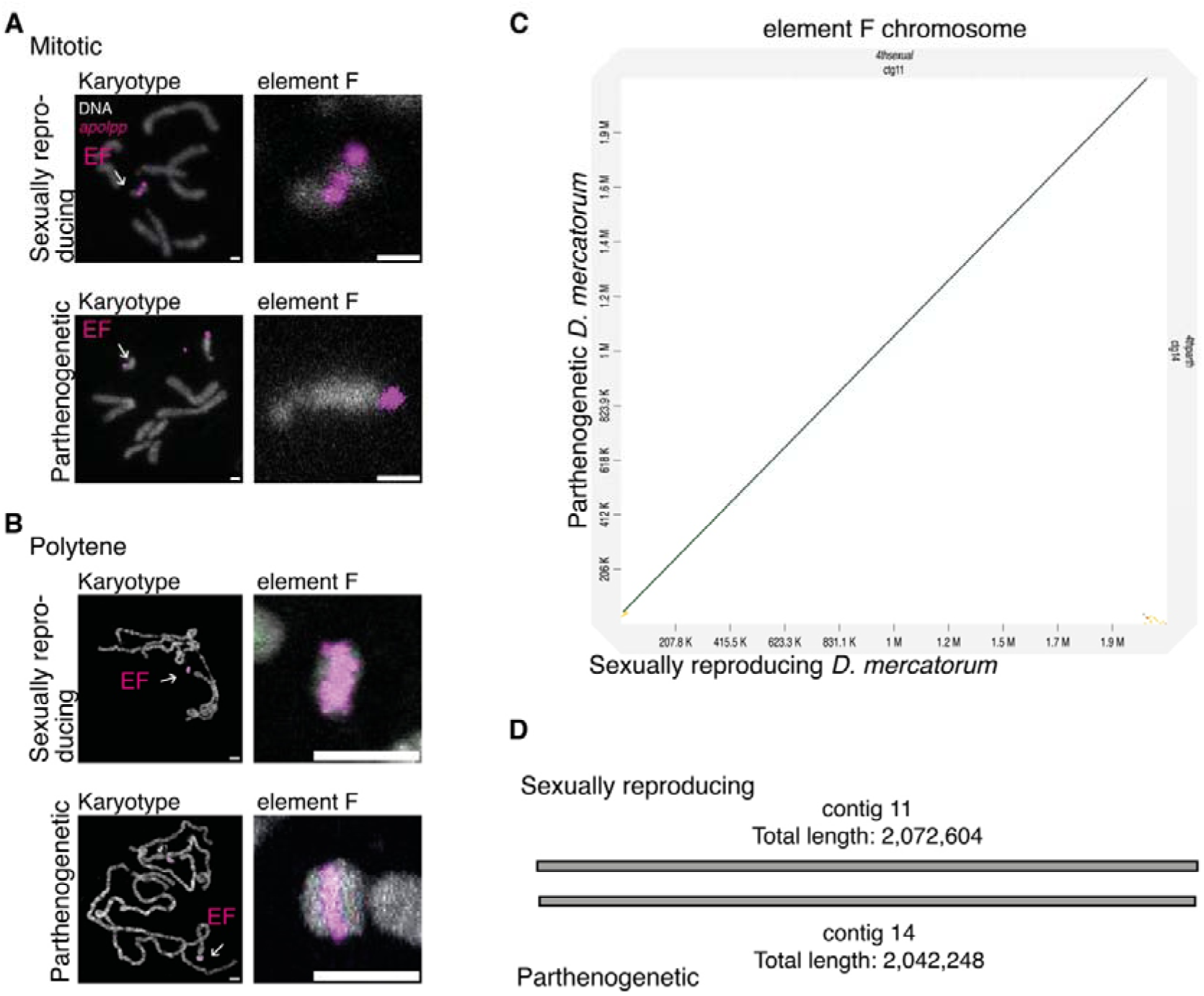
Element F chromosome differences between sexually reproducing and parthenogenetic *D. mercatorum*. **A**) Mitotic chromosome spread from the larval brain of parthenogenetic and sexually reproducing *D. mercatorum* 3^rd^ instar larvae. *In situ* to *apolpp* (magenta). The DNA is in monochrome. Scale bar, 10 μm. **B**) Polytene chromosome spread from the salivary glands of parthenogenetic and sexually reproducing *D. mercatorum* 3^rd^ instar larvae. *In situ* to *apolpp* (magenta). The DNA (DAPI) is in monochrome. The scale is 10 μm. **C**) Alignment of the coding region of the parthenogenetic against the sexually reproducing *D. mercatorum* element F. The non-coding region is not assembled. **D**) Schematic of the matching contigs representing the coding region of element F.

As the sexually reproducing strain of *D. mercatorum* and the parthenogenetic strain originate from different geographical locations, we asked whether the changes in chromosomal organisation between these strains could also be seen in other geographical isolates. To this end, we examined 18 strains of *D. mercatorum* strains and determined whether they were sexually reproducing, facultative parthenogenetic, or fully parthenogenetic. The six sexually reproducing strains could generate embryos parthenogenetically but these did not develop into adult offspring (Fig. S3). The eight facultative parthenogenetic strains were sexually reproducing strains capable of producing parthenogenetic offspring or previously categorized as such by the National Drosophila Species Stock Center or earlier studies (Sperling, et al. 2023). Finally, the four parthenogenetic strains could be maintained as fully female-only stocks (Fig. 4A-C, Data Table S1O).

**Figure 4:**
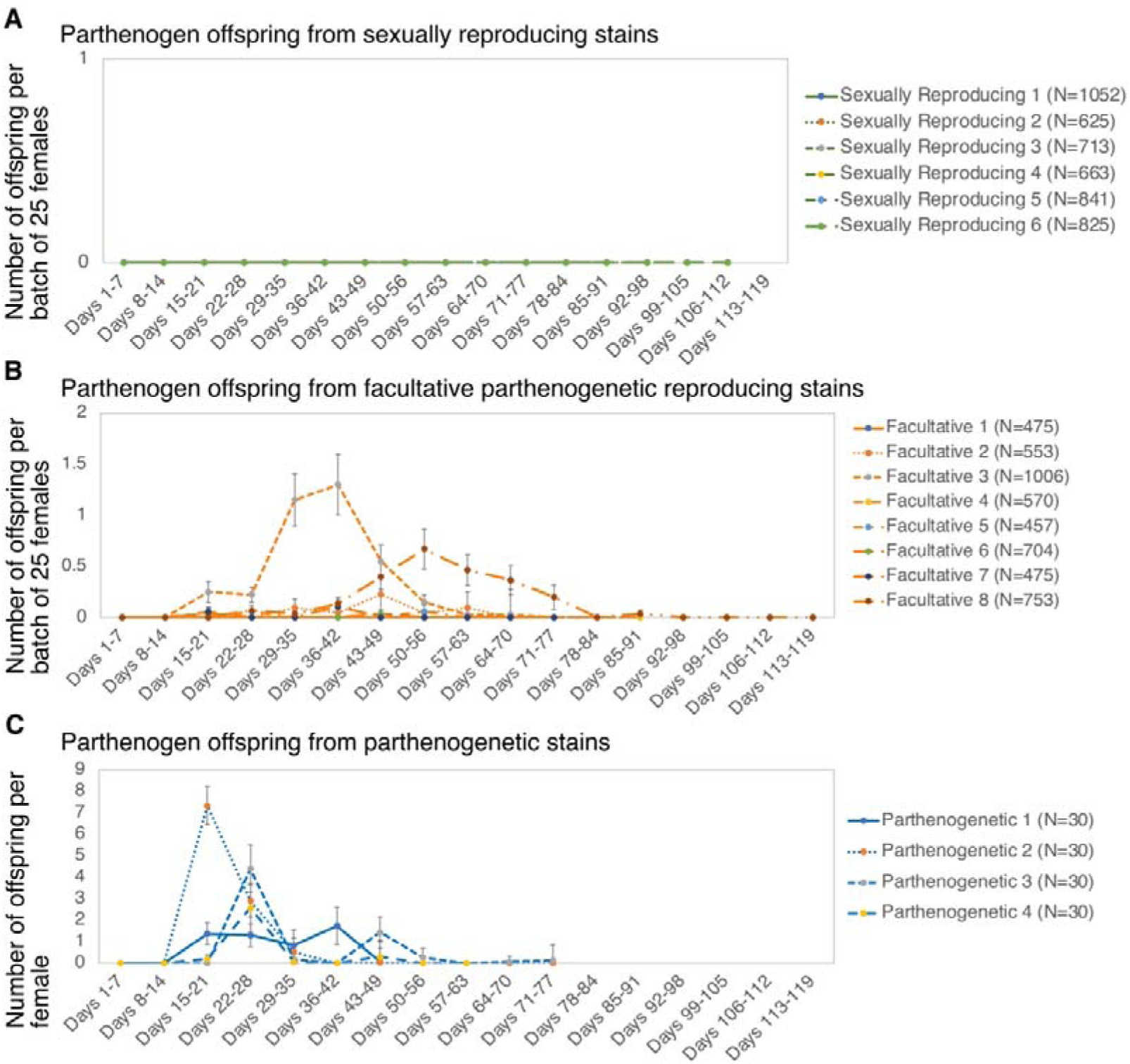
Parthenogenetic ability of *D. mercatorum*. **A-C)** Plot of parthenogenetic offspring per batch of 25 virgin females for sexually reproducing and facultative parthenogenetic strains, and per single virgin female for parthenogenetic strains. The error bars represent standard error of the mean.

Of strains with known geographical origins, the fully parthenogenetic strains and the facultative parthenogens had been collected from Hawaii and Brazil. In contrast, virgin females of strains from other locations in North America, South America, and Africa did not produce adult offspring in these assays (Fig. 5). We found that only two of the Brazilian strains did not have all three inversions on chromosomal element B. These included the original sexually reproducing stock, which lacked all three inversions (Fig. 5, Fig. 2A; Fig. S4-6) and another sexually reproducing stock that had two of the three inversions but lacked the most distal one, *In(EB)A* (Fig. 5, Fig. S4D). The other *D. mercatorum* strains had all three inversions (Fig. 5 and Fig. S4-6). Thus, all parthenogenetic, facultative, and sexually reproducing strains, except for one sexually reproducing strain, carried the *In(EB)B* inversion. The fully parthenogenetic strain that produced the most offspring per female also carries two large insertions (Fig. 5A, S6B). These insertions may bring increased fitness as this strain is the healthiest female-only stock and produces the highest number of parthenogenetic offspring (Data Table S1O). The presence of the three large element B inversions in all strains outside Brazil and their presence in both parthenogenetic most sexually reproducing strains suggests that this inversion may also contribute to overall fitness and that it likely arose before strains became parthenogenetic (see Discussion).

**Figure 5:**
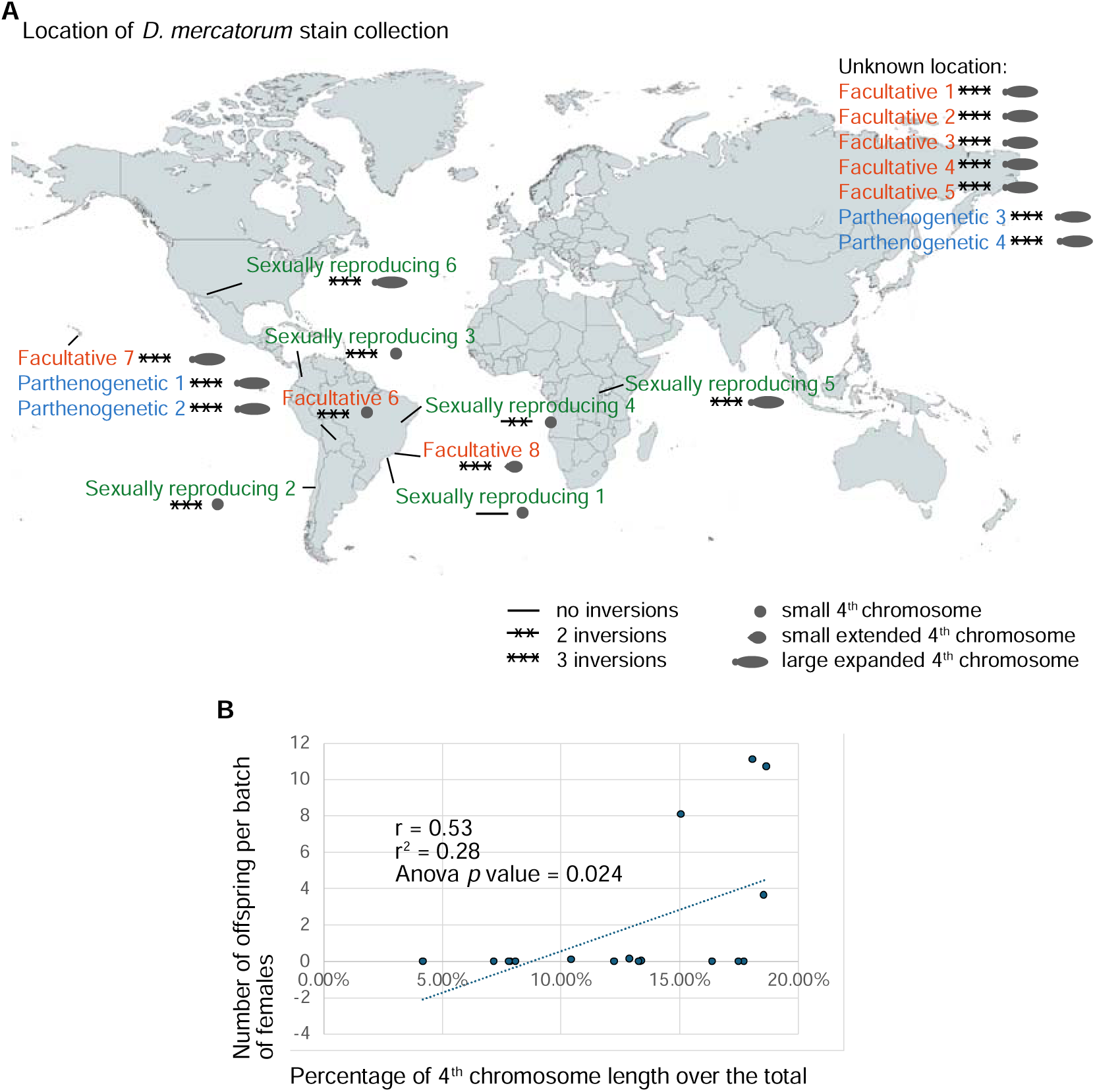
Geographical collection points of the different strains of *D. mercatorum* with the genomic differences indicated. **A**) World map indicating the parthenogenetic capability of the strains from their collection location, the inversions present on element B and the length of element F. The map was rendered using MapChart. Of the sexually reproducing strains from Brazil, one (our reference strain) had no inversions in element B and no expansion of element F, whereas the other had two of the three inversions we previously characterized for element B; neither showed any expansion of element F. Four other sexually reproducing strains all showed the three inversions of Element B with two of them having a dot-like and two, an expanded element F. There is no clear correlation of these chromosomal variants with their widespread geographical distribution suggesting that they arose some time ago, possibly in Brazil, where the changes were less extensive, and then dispersed. Strains that undertake sexual reproduction appear to have the minimal genome reorganization (zero or only 2 inversions) and originate in Brazil. Of the facultatively parthenogenetic strains, all had the three inversions with 6 having an expansion of element F, one having partial expansion of element F and one with a dot-like element F. **B)** Plot of the element F length as a percentage of the total chromosome length, compared to the number of parthenogenetic offspring per batch of 25 sexually reproducing or facultative females, and per single female for the fully parthenogenetic strains. The *p*-value was calculated using an ANOVA test.

Our examination of the element F chromosome in all 18 strains confirmed it to be of uniform size in polytene chromosomes whereas it varied in size in the mitotic chromosomes of larval neuroblasts (Fig. 3A,B, S7-S8). This is in line with the expanded part of the chromosomal element being heterochromatic, likely as a result of the expansion of simple sequence DNA. The strains with the elongated element F were collected outside of South America (Fig. 5A). All facultative and fully parthenogenetic strains carried an expanded element F chromosome, except for one strain with a small element F that showed a very low level of facultative parthenogenesis. The length of the element F chromosome positively correlated (n = 18, Pearson coefficient 0.53, ANOVA *p*-value = 0.024) with increased parthenogenetic ability (Fig. 5B). The finding that most facultative and all fully parthenogenetic strains possess increased heterochromatin on the F chromosomal element suggests that this particular aspect of chromosomal organization may contribute directly or indirectly to parthenogenetic reproduction.

## Discussion

Here we examine the thesis that large scale changes in genome organization accompany, and thereby possibly facilitate, the transition from sexual to parthenogenetic reproduction. To this end we have compared the genomes of geographically separated, sexually and parthenogenetically reproducing strains of *D. mercatorum* to evaluate the nature and spread of large-scale genomic changes to chromosomal elements B and F that have been previously identified (Wharton 1943; Sperling, et al. 2023). Our previous comparison single strains of sexually reproducing, facultative parthenogenetic, and parthenogenetic strains of *D.mercatorum* identified three genes, *Desat2*, *polo*, and *Myc*, as causal for parthenogenesis in *D. mercatorum.* Accordingly, we were able to recapitulate the changed expression levels of these three genes in *D. melanogaster* and so drive parthenogenesis in this otherwise non-parthenogenetic species. Since the expression of none of these three genes was linked to the larger genomic changes, we sought to understand whether the differences in organization of the sexually and parthenogenetically reproducing *D. mercatorum* genomes were connected to parthenogenetic reproduction and/or geographical adaptation. We were unable to determine any strong correlation of the occurrence of the inversions on element B and expansion of heterochromatin on element F with geographical collection locations beyond migration out of Brazil. However, F-element expansion did correlate with the ability to reproduce parthenogenetically, leading us to propose a model for a possible sequence of evolutionary events (Fig. 6). It seems likely that the generation of first two, then three inversions on element B precedes the transition to parthenogenetic reproduction that in turn would precede expansion of the F element because the great majority of such flies that have a small dot-like element F reproduce sexually. Conversely, this expansion of element F leads to a dramatic increase in parthenogenesis. Of eleven strains able to reproduce by either full or facultative parthenogenesis, only one had a dot-like element F and that strain showed very poor facultative parthenogenesis.

**Figure 6.**
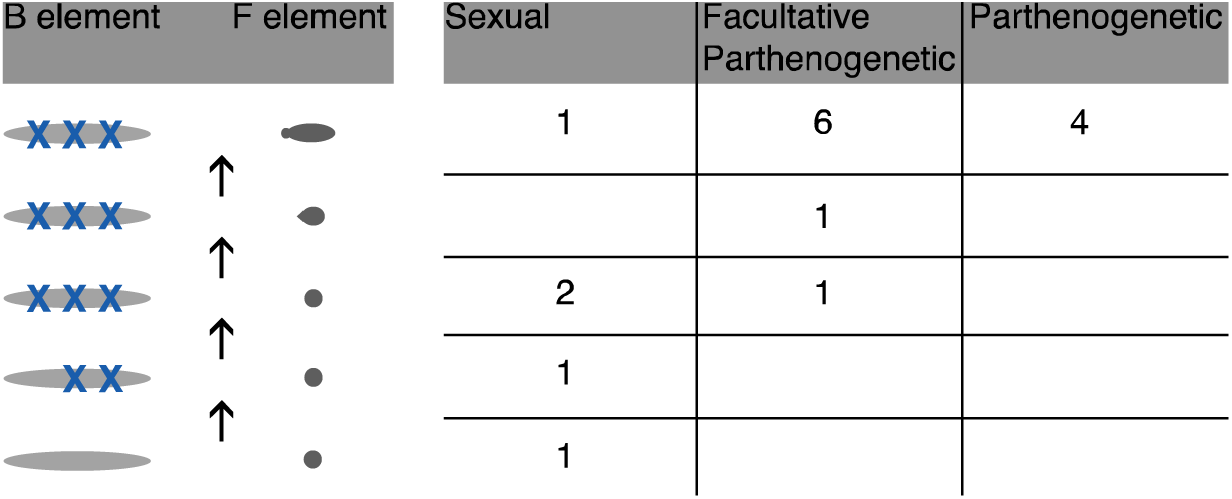
Model for changes in genome organisation that accompany parthenogenesis. Inversions on the B element are indicated by the “X” symbol and are indicated to arise successively although alternative models are possible. The expansion of element F from its dot-like origins are also depicted. The table to the tight indicates the numbers of strains from the 18 examined that fall into the different categories of chromosomal arrangements.

We might speculate that the chromosome inversions we observe may drive the development of unique characteristics that facilitate the subsequent development of parthenogenesis. However, we have no evidence, one way or another, that these inversions change gene expression in a manner necessary for parthenogenesis. Chromosome inversions have the potential to lead to speciation, as has been suggested in the repleta group species of *Drosophila,* of which *D. mercatorum* is a member (Wasserman 1960; O’Grady and Markow 2012). The repleta species group consists of cactophiles, thought to originate from South America (O’Grady and Markow 2012; Acurio 2024) but there are strains that have evolved to be cosmopolitan and feed on human food waste in urban environments (Wasserman 1960; O’Grady and Markow 2012). The presence of the three inversions we characterize in widely distributed strains, predominantly from urban environments, may reflect an expansion beyond their original range due to an ability to accommodate a diversified diet (Wasserman 1960). We identified four genes in the vicinity of the breakpoints of these inversions that are differentially expressed in the ovaries of sexually compared to parthenogenetically reproducing flies. We speculate that the changes in expression of these genes might be linked to an ability to enhance parthenogenesis, but we have not tested this in *D. melanogaster* as we did previously for *Desat2*, *polo*, and *Myc* (Sperling et al., 2023). This is because we have not, as yet, been able to further increase the number of genetic changes that lead to parthenogenesis in *D. melanogaster* without observing a decrease in parthenogenesis (Data Table S1L). This is likely a consequence of a decrease in fitness, as it was difficult to generate the flies carrying more than three alleles. Unfortunately, in our hands, *D. mercatorum* is not tractable for transgenesis, and so we need further work to improve our system for manipulating transgenes in *D. melanogaster* in order to test these present findings. It is also possible that the effect of the chromosome inversions could be indirect, since chromosome inversions suppress recombination when heterozygous. This could maintain the co-inheritance of gene clusters and so contribute to the maintenance of isolation between different populations, such as parthenogenetic versus sexually reproducing strains or cosmopolitan versus rural strains of *D. mercatorum*. Indeed, it has been proposed that linkage disequilibrium between alleles or decreased recombination near inversions could be selected for in migrating populations if they contain loci that are involved in local adaptation (van Delden and Kamping 1991; Veuille, et al. 1998; Andolfatto, et al. 1999; Kirkpatrick and Barton 2006). It is also described that inversions can be fixed in an isolated geographical location due to a colonization event and thus not likely due to adaptation (Westram, et al. 2022). However, we note that the inversions we observe in element B of *D. mercatorum* strains are geographically widespread and so if they have evolved through isolation, they are not constrained by it.

The expansion of the element F chromosome, which correlates strongly with ability to reproduce parthenogenetically, is also connected to the geographical collection point since it is present in all strains collected outside of South America. Heterochromatin is largely composed of repetitive sequences, and in the F-element of *D. melanogaster,* chromosome 4, it comprises simple sequence repeats and an accumulation of transposable elements. The repetitive nature of such sequences makes it difficult to assemble them into contigs in whole genome sequencing. Future experiments could reassemble the genome with Hi-C genome sequencing data in order to resolve more of these repetitive sequences. The heterochromatin changes in *D. mercatorum* can also be compared to those of other *Drosophila* species or strains with expanded element F chromosome arms. These include the facultatively parthenogenetic *Drosophila ananassae* (Matsuda and Tobari 2004), in which the expanded sequence is composed of retrotransposons (Leung, et al. 2017), and strains of several other species collected in Hawaii where their expanded sequence appears to be composed of simple sequence, satellite DNA (Craddock, et al. 2016).

In *D. melanogaster,* the effect of a heterochromatic environment on gene expression has been intensively studied. Classical genetic observations by Muller first described the phenomenon known as position effect variegation (PEV), whereby translocation of a gene from a euchromatic to a heterochromatic environment stochastically alters gene expression levels (Muller, 1930), which become fixed in a cellular lineage leading to a variegated pattern. Reciprocally, it was also shown that *cubitus interruptus,* a gene located in the heterochromatic environment of chromosome 4, exhibited PEV when translocated to a euchromatic region. Although classically studied through chromosomal rearrangements, recent studies of PEV of *white* transgenes inserted into multiple sites on chromosome 4 of *D. melanogaster*, have permitted investigation of the arrangement of heterochromatin on this tiny chromosome, which is marked by a chromosome 4-specific chromosomal protein (POF (painting-of-fourth), and utilizes a dedicated histone methyltransferase, EGG (reviewed in Riddle and Elgin, 2006; Riddle et al., 2009). In several respects, facultative parthenogenesis resembles a form of variegated gene expression with the added complexity that parthenogenesis requires multifactorial events. This leads us to speculate that the onset of parthenogenesis in *D. mercatorum* could be promoted by an expansion of heterochromatin on element 4 resulting in changes in gene expression resembling those brought about through PEV. Such changes might then re-adjust over multiple fly generations to stabilise gene expression in the new expanded heterochromatic environment. A comparison of the parthenogenetic and sexually reproducing transcriptome datasets indicates that there are five differentially expressed genes present on element F, namely *apolpp*, *CG33978*, *Thd1*, *NfI*, and *mGluR*. Notably, the transcription factor encoded by *NfI* physically interacts with the Polycomb group (PcG) protein encoded by *Sex comb on midleg* (*Scm*) (Shokri et al., 2019), which in turn physically interacts with Suppressor of variegation 3-3 (Su(var)3-3) that promotes heterochromatin formation (Kang et al., 2015). Further studies would be required to determine whether there is a link between *NfI* expression as a result of the expansion of heterochromatin we observe on chromosome element F and the origins of parthenogenesis in *D. mercatorum*.

There is more to the biology of parthenogenesis than the change(s) that enable the initiation of development of an unfertilized egg. Some genomic changes may arise before parthenogenesis, predisposing a species to establish a parthenogenetic lineage. Other changes might arise after a minimal set of events enabling parthenogenesis to have occurred, either as a direct or indirect consequence of the genetic changes enabling parthenogenetic reproduction. Untangling the temporal progression of these events will be challenging. Our present study suggests that the element B inversions seen in parthenogenetic populations may have enabled this species to spread, whereas the expansion of heterochromatin in element F positively correlates with parthenogenetic ability. The next challenge will be to understand the connection between the effects of element F heterochromatin upon parthenogenesis to determine exactly how these two phenomena are related.

## Materials and Methods

### Drosophila strains

All *Drosophila* strains used in this study are listed in Data Table S1L and S1O.

### *D. mercatorum* genomes and transcriptomes

The previously published *D. mercatorum* genomics and transcriptomics: Alexis L Sperling, Daniel K Fabian, Erik Garrison, and David M Glover, 2023, Genetic basis for facultative parthenogenesis in *Drosophila*, ENA (European Nucleotide Archive), PRJEB43100. The data analysis is described in detail also on https://github.com/ekg/drosophila and https://github.com/FabianDK/Dmerc and in Sperling et al. (2024) (Sperling, et al. 2024). Additional analysis was carried out with MUMmer4 (Marcais, et al. 2018). Numer was used to create a delta file which was then used to quantify the difference between the whole parthenogenetic and sexually *reproducing D. mercatorum* genomes. Subsequently, careful examination revealed the exact locations of the larger inversions on element B.

## Alignment

The genome alignment was performed with Minimap2 v2.24 through D-genies (Cabanettes and Klopp 2018). This generated differing results from our previous analysis (Sperling, et al. 2023) and from using Mashmap v2.0 also through D-genies (Cabanettes and Klopp 2018). Of these analyses, only Minimap2 v2.24 alignment gave the same results as the physical mapping when the input sequence was provided in the correct direction.

## Inversion mapping

The inversions on element B were identified using MUMmer4: first using nucmer to create the delta file and then using the show-coords with –d –l options. The output file shows the mapping of segments of the genome and the inversions are easily identified in the [FRM] column, after <10kb repetitive regions are discounted. The locations of the inversions were then searched in a basic local alignment search file (BLAST) 2.9.0+ (Zhang, et al. 2000) for element B. The genes near the inversion points were then identified from the DNA sequence and the sequences were taken from IGV and back verified against both parthenogenetic and sexually reproducing *D. mercatorum* genomes using BLAST 2.9.0+.

## Manual gene annotation curation

Manual curation of the gene annotation was performed on the genes in the 100Kbp region flanking each of the paracentric inversions on element B and the genes within each of the 3 identified inversions. This was performed using nucleotide BLAST against the *D. melanogaster* (release 6) reference genome and against both parthenogenetic and sexually reproducing *D. mercatorum* genomes to eliminate the possibility of gene duplications. The genome was then walked using IGV and the annotation was manually checked for each gene.

## Mitotic chromosome preparation

*D. mercatorum* brains were dissected from 3^rd^ instar larvae in saline (0.7%NaCl) and cultured for 1 h in 10 μM colchicine in 0.7% NaCl. The brains were then subjected to hypotonic shock by incubation in 0.5% trisodium citrate for 9 min and fixed in a 60 sec incubation in 45% acetic acid followed by 5 min in 60% acetic acid on a coverslip. A slide was placed over the coverslip and squashed between two sheets of blotting paper. Immediately after squashing, the preparation was frozen in liquid Nitrogen and the coverslip removed using a scalpel. The squashed brain preparation was then washed for 5 min in 70% and 5 min in 100% ethanol then air-dried.

If an *in situ* was to be performed, the slides were baked at 58°C for 1 h in a dry oven. The preparations were then denatured with 70% formamide in 2×SSC at 70°C for 20 min. The brain preparation was then washed for 5 min in 70% and 5 min in 100% ethanol then air-dried before proceeding onto the HRC protocol.

## Polytene chromosome preparation

Salivary glands were dissected from 3^rd^ instar larvae and prepared for *in situ* hybridization using a protocol adapted from the Duronio Lab (https://theduroniolab.web.unc.edu/resources#protocols). Place 6 ul 1:2:3 lactic acid:water:acetic acid on a cover slip. Dissect salivary gland in 0.7% NaCl. Fix gland for 30-60 seconds in 45% acetic acid, Transfer gland to drop of 1:2:3 on cover slip. Cover with slide and squish. Dip slide in liquid nitrogen until bubbling stops. Breath on frozen slide to get frost and flip off cover slip with a razor blade. Wash 3 x 10 min in 95% ethanol and allow to air dry. Denature Chromosomes by incubate slides in 2 x SSC (saline sodium citrate buffer) at 65°C for 30 min. Place in 2 x SSC at RT for 10 min. Incubate in 70 mM NaOH for 3 min. Rinse in 2 x SSC. Soak slides in 70% ethanol 2 x 5 min and in 95% ethanol 2 x 5 min

## Molecular Instruments HCR sample preparation

All materials including buffers and probes were purchased or gifted from Molecular Instruments. To ensure optimal hybridization to mitotic chromosome preparations. The standard Molecular Instruments HCR protocol (v3.0) for sample on slide was used (Choi, et al. 2018).

## Imaging

All images were acquired on a Leica SP8 confocal microscope, and the images were minimally optimized for brightness and contrast using ImageJ/FIJI (Schindelin, et al. 2012). Chromosome length measurements were performed in FIJI. No other image alteration was performed.

## Statistical analysis

The Chi Squared Test (in R) was used to determine if the changes in gene expression within the inverted regions were significant. A T test (in R) was used to determine if the differences in element F chromosome length were significant. An ANOVA test was used to determine if there was a correlation between parthenogenetic ability and element F chromosome length.

## Facultative Parthenogenesis assay

The facultative parthenogenesis assay was performed as previously published (Sperling and Glover 2023b).

## Data Availability

All data used in this study were previously published and are publicly available: Alexis L Sperling, Daniel K Fabian, Erik Garrison, and David M Glover, 2023, Genetic basis for facultative parthenogenesis in *Drosophila*, ENA (European Nucleotide Archive), PRJEB43100.

## Acknowledgements

We would like to thank Paula Almeida-Coelho for technical advice on in situs and karyotyping. We are grateful to the Cambridge Genetics Fly Facility for their support of this work.

## Supplementary Data

Data tables are in a separate file.

**Data Table 1: Manual annotation of the genes within *In(EB)A*, *In(EB)B*., and *In(EB)C*; differentially expressed genes; element F annotation and differentially expressed genes sexually reproducing and parthenogenetic strains of *D. mercatorum*; and parthenogenesis assay. A**) The transcripts mapped within the whole *In(EB)A* inversion. **B**) The genes 100kb of breakpoint 1. **C**) The genes 100kb of breakpoint 2. **D**) The transcripts mapped within the whole *In(EB)B* inversion. **E**) The genes 100kb of breakpoint 3. **F**) The genes 100kb of breakpoint 4. **G**) The transcripts mapped within the whole *In(EB)C* inversion. **H**) The genes 100kb of breakpoint 5. **I**) The genes 100kb of breakpoint 6. **J**) The transcripts mapped for the entire genome. **K**) Differentially expressed genes on element B between the mature eggs of sexually reproducing and parthenogenetic strains of *D. mercatorum*. **L**) *Drosophila melanogaster* parthenogenesis assay for three alleles including gene name, genotype, culture temperature, number of batches, total number of females tested, average max lifespan, average maternal age of parthenogenesis, proportion of total lifespan when parthenogenesis occurs, number of offspring, percent of offspring, stock number. **M**) Manual annotation of the genes on element F. **N**) Differentially expressed genes on element B between the mature eggs of sexually reproducing and parthenogenetic strains of *D. mercatorum*. **O**) *Drosophila mercatorum* parthenogenesis assay including name, species, genotype, culture temperature, number of batches, total number of females tested, average max lifespan, average maternal age of parthenogenesis, proportion of total lifespan when parthenogenesis occurs, number of adult offspring, percent of adult offspring, offspring information, stock origin, and stock number.

**Figure S1:**
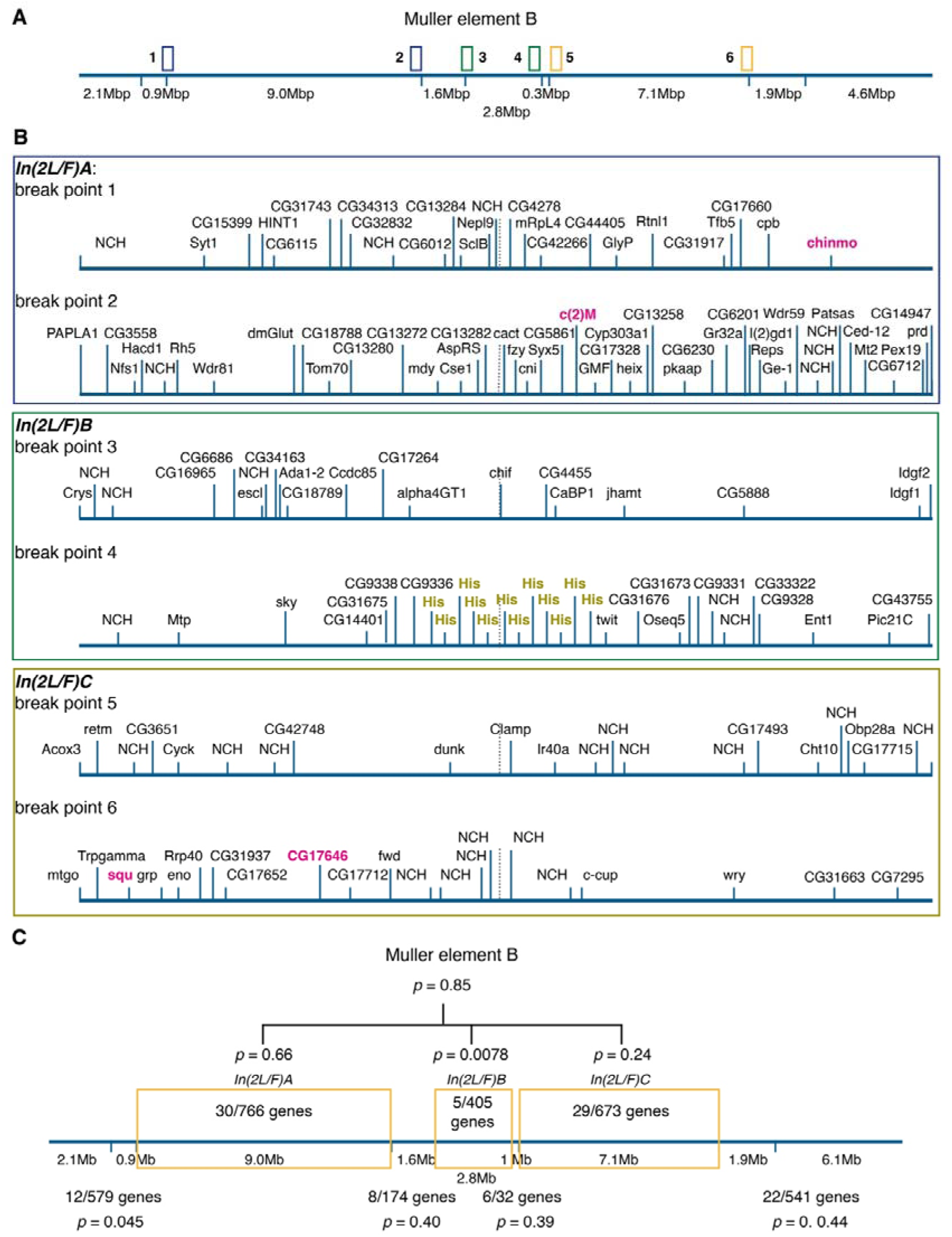
Genes within 100kb from the break points of the inversions present on element B. **A**) Schematic of the breakpoints. **B**) Genes present at the specific breakpoints with the differentially expressed ones indicated in pink. Annotated genes that have no close homology (NCH) are indicated as such. **C**) Analysis for the enrichment of differentially expressed genes within the element B inversions. The comparison is between the specific inversion compared to the not inverted segments in the rest of the chromosome arm. The p values were calculated using the Chi squared test.

**Figure S2:**
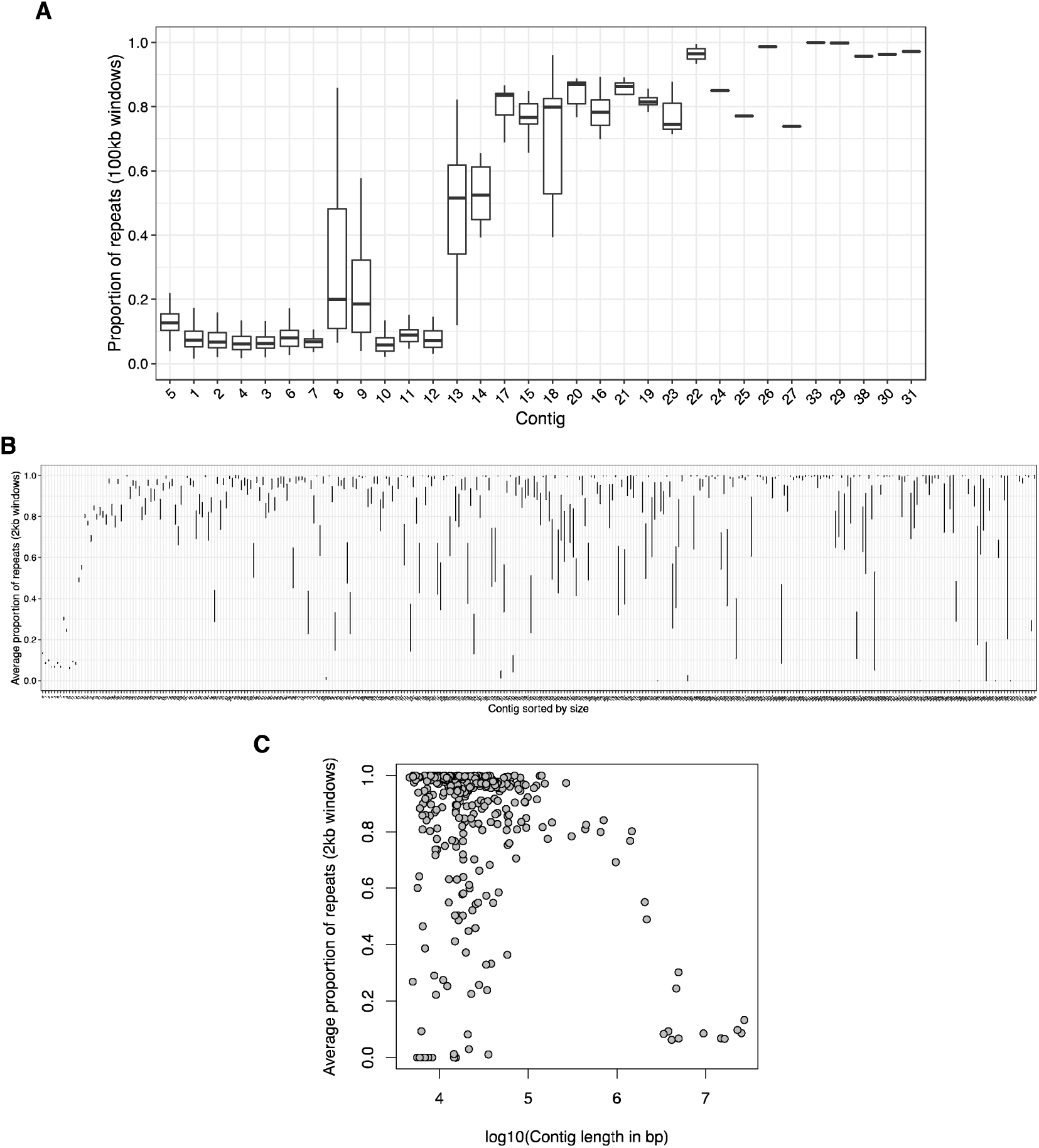
Repetitive content of the parthenogenetic genome. **A**) The average proportion of repeats (per 100Kb window) plotted for each contig from the largest to the smallest showing that the smaller contigs were largely composed of repetitive DNA. **A**) The average proportion of repeats (per 2Kb window) plotted for each contig from the largest to the smallest showing that the smaller contigs were largely composed of repetitive DNA. **B**) The average proportion of repeats (per 2Kb window) plotted against the log_10_ of the contig length.

**Figure S3:**
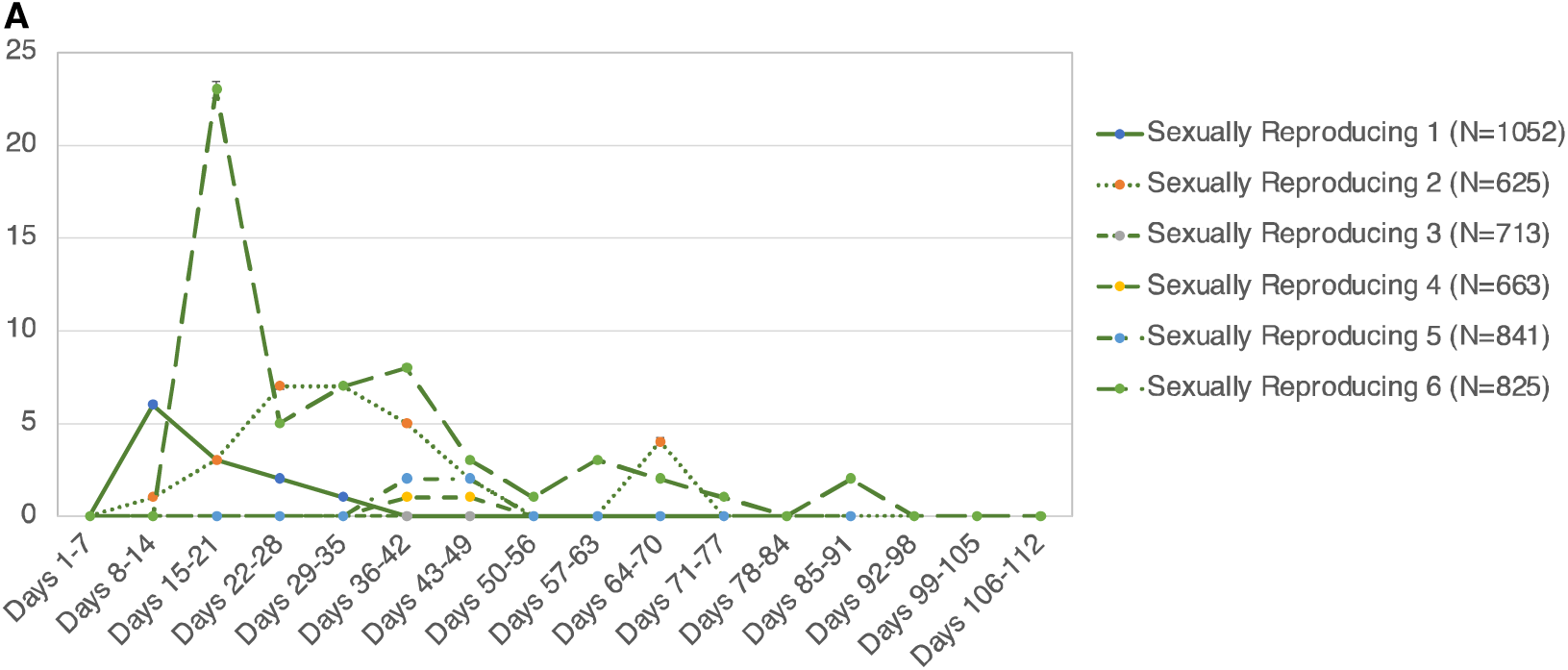
Parthenogenetic ability of *D. mercatorum*. **A)** Plot of parthenogenetic embryos per batch of 25 virgin female flies for sexually reproducing strains. The error bars represent standard error.

**Figure S4:**
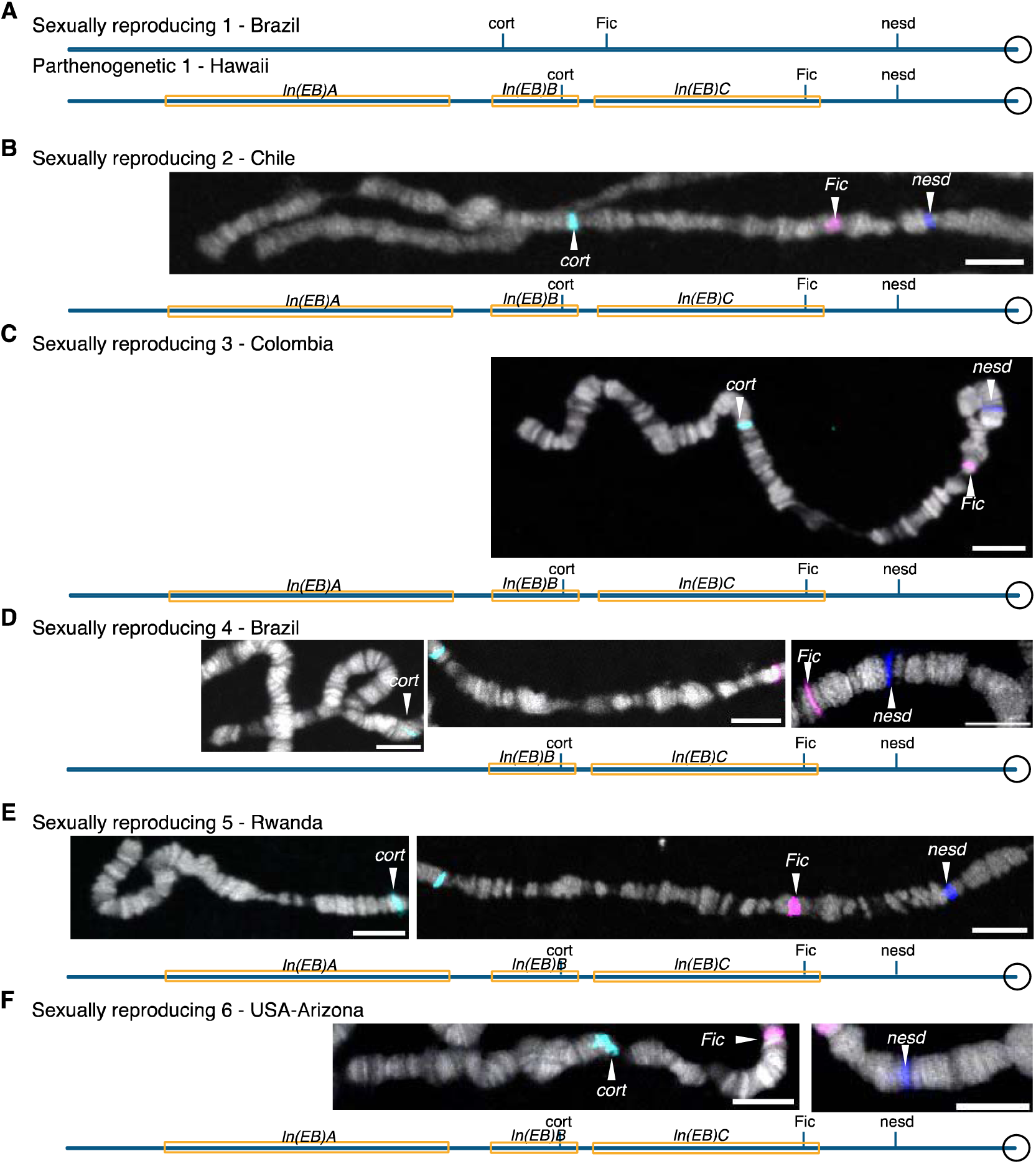
Polytene element B chromosomes of sexually reproducing salivary glands of 3^rd^ instar *D. mercatorum* larvae. **A**) Layout of the Sexually reproducing 1 and Parthenogenetic 1 element B polytene chromosomes. **B-F**) Polytene chromosome from Sexually reproducing 2-6. *In situs* to select genes on the largest contigs to *cort* (cyan), *Fic* (magenta), and *nesd* (blue). The DNA (DAPI) is in white. The scale is 10 μm. A schematic of element B is given below the images.

**Figure S5:**
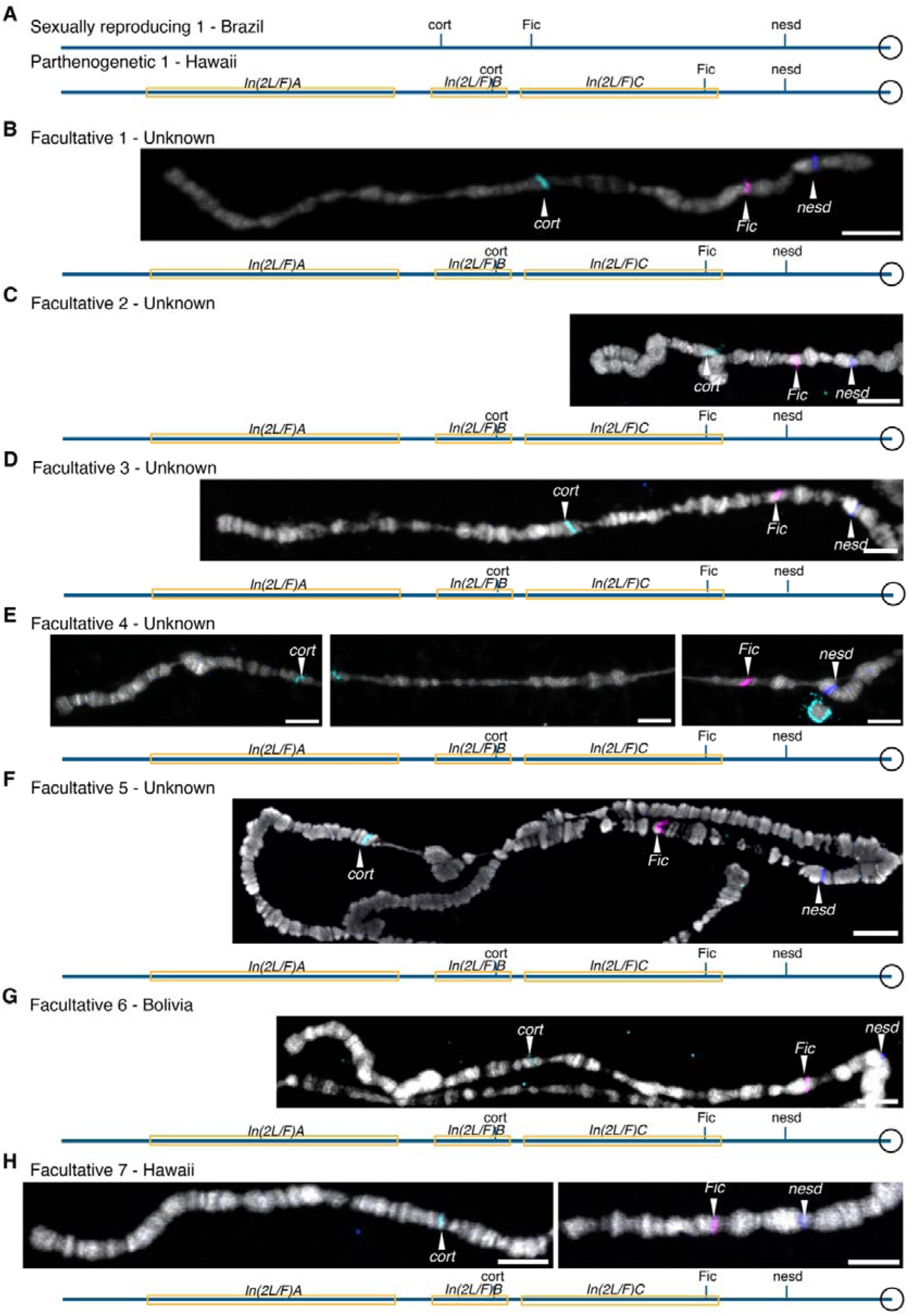
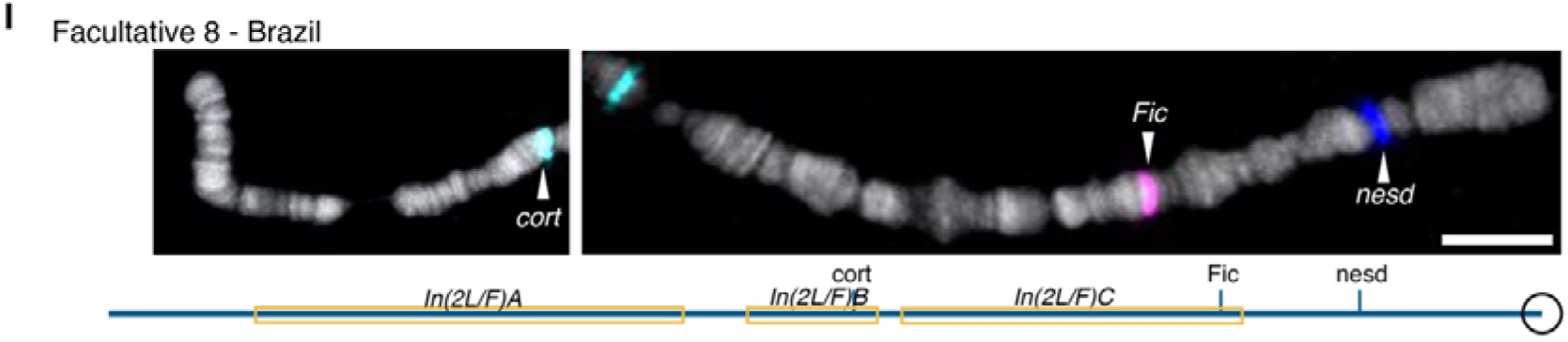
Polytene element B of facultative parthenogenetic reproducing salivary glands of 3^rd^ instar *D. mercatorum* larvae. **A**) Layout of the Sexually reproducing 1 and Parthenogenetic 1 element B polytene chromosome. **B-I**) Polytene chromosome from Facultative parthenogenetic 2-8. *In situs* to select genes on the largest contigs to *cort* (cyan), *Fic* (magenta), and *nesd* (blue). The DNA (DAPI) is in white. The scale is 10 μm. A schematic of the element B is given below the images.

**Figure S6:**
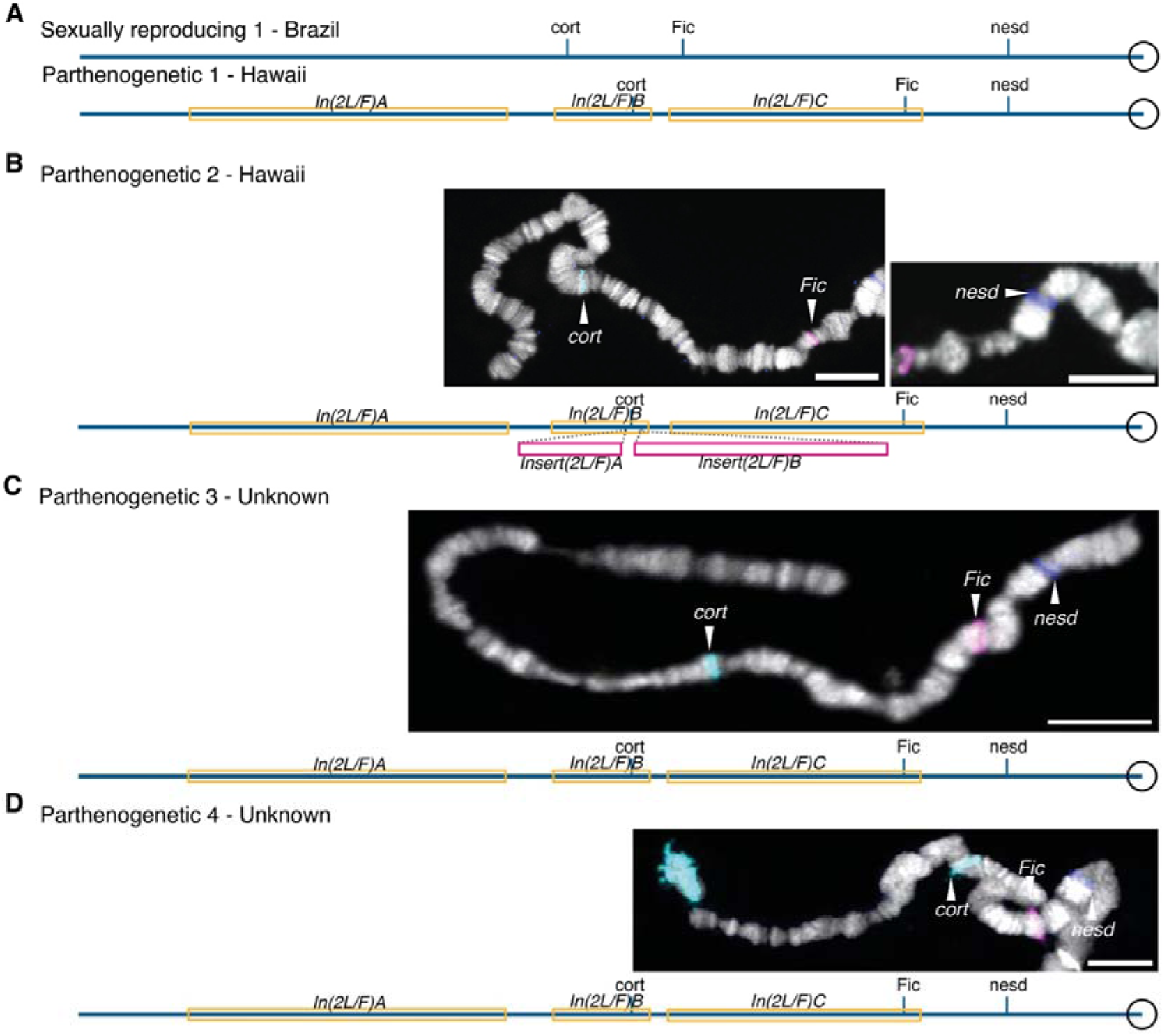
Polytene element B chromosomes of Parthenogenetic reproducing salivary glands of 3^rd^ instar *D. mercatorum* larvae. **A**) Layout of element B polytene chromosome for Sexually reproducing 1 and Parthenogenetic strains 1. **B-D**) Polytene chromosome from Parthenogenetic 2-4. *In situs* to select genes on the largest contigs to *cort* (cyan), *Fic* (magenta), and *nesd* (blue). The DNA (DAPI) is in white. The scale is 10 μm. A schematic of element B is given below the images.

**Figure S7:**
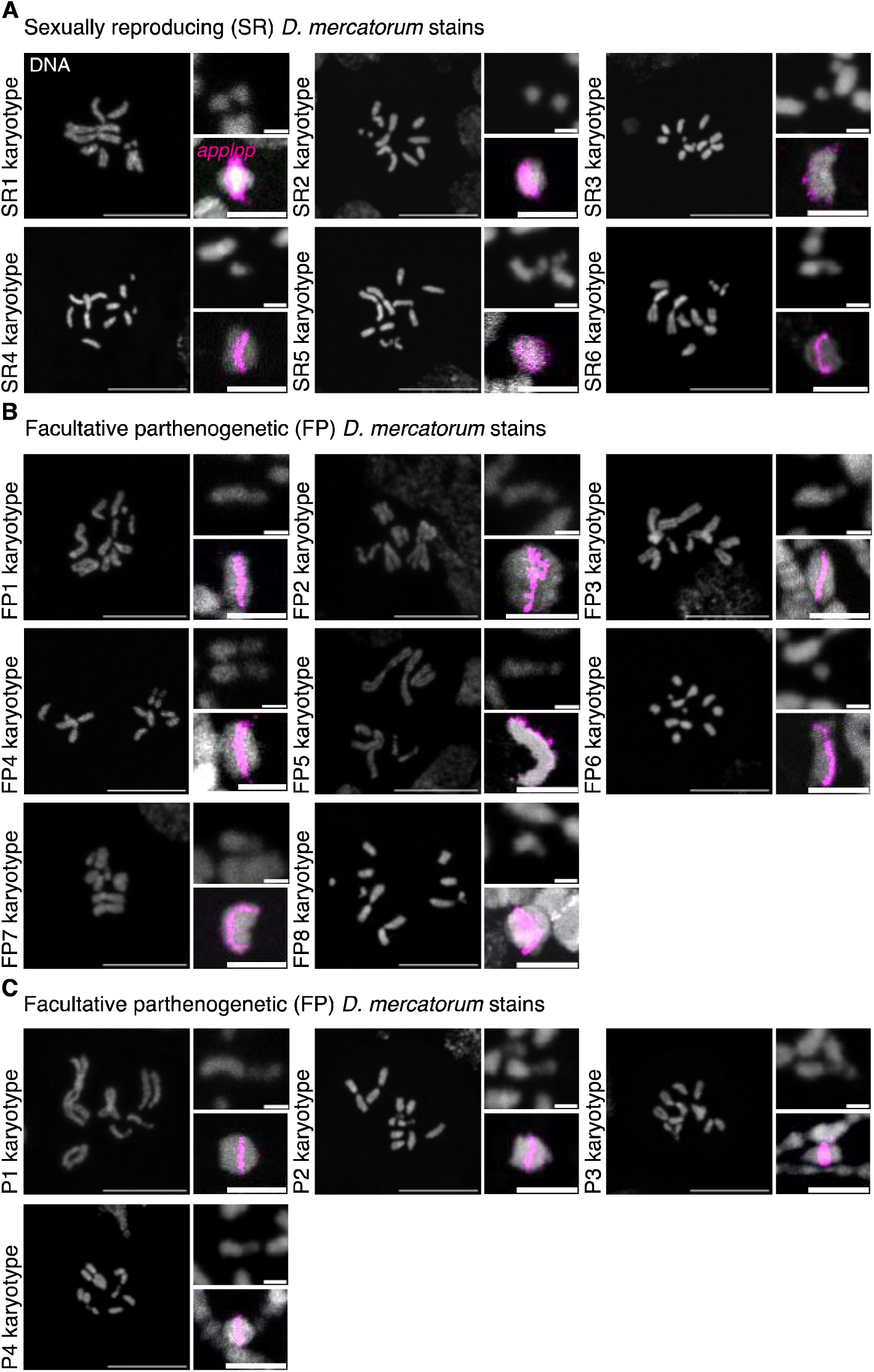
Mitotic karyotypes of 18 *D. mercatorum* stains and polytene element F. **A**) Sexually reproducing (SR) 1-6 karyotypes with magnified images of element F and the polytene element F for size comparison. **B**) Facultative parthenogenetic (FP) 1-8 karyotypes with magnified images of element F and the polytene element F for size comparison. **C**) Parthenogenetic (P) 1-4 karyotypes with magnified images of the element F and the polytene element F for size comparison. *In situ* to *apolpp* (magenta) on the polytene chromosome arms. The DNA (DAPI) is in white. The scale is 10 μm and 1 μm in the magnified images.

**Figure S8:**
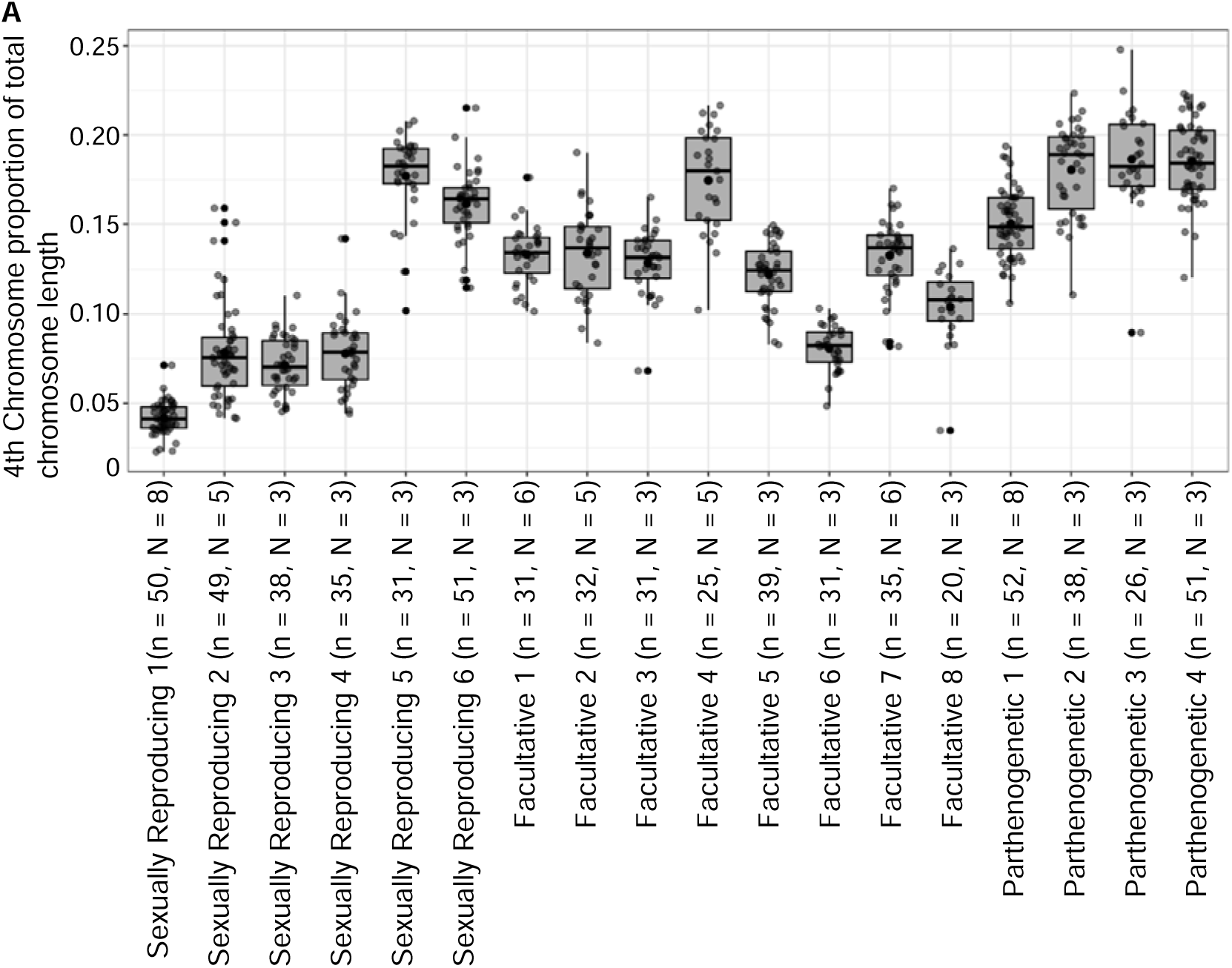
Element F chromosome length for 18 different strains of *D. mercatorum*. **D**) Box plot of element F chromosome length measured in confocal images using FIJI, normalized over the proportion of total chromosome length. The data for Parthenogenetic 1 and Sexually Reproducing 1 are also in Figure 1D.

